# Decoding the genetic and chemical basis of sexual attractiveness in parasitic wasps

**DOI:** 10.1101/2023.01.09.523239

**Authors:** Weizhao Sun, Michelle Ina Lange, Jürgen Gadau, Jan Buellesbach

**Affiliations:** Institute for Evolution & Biodiversity, University of Münster, Hüfferstr. 1, DE-48149 Münster, Germany

**Author notes:** **Author Contributions:** J.B. conceived the study; J.B. & W.S. designed the study; W.S., & M.I.L. performed the experiments and collected the data, W.S., J.B. & M.I.L. analyzed the results; W.S. & J.B. validated the results, W.S. & J.B. visualized the data; J.B. & J.G. provided the resources; J.B. & W.S. wrote the original draft; J.B., W.S. & J.G. reviewed and edited the final manuscript version, J.B. & J.G. supervised the project; J.B. acquired the funding for the study.

**Keywords:** Sex pheromones, cuticular hydrocarbons, fatty acid synthase, chemical communication, Hymenoptera

## Abstract

Attracting and securing potential mating partners is of fundamental importance for successfully initiating reproduction and thus assuring the passing of genes to the next generation. Therefore, signaling sexual attractiveness is expected to be tightly coordinated in communication systems synchronizing senders and receivers. Chemical signaling has permeated through all taxa of life as the earliest and most wide-spread form of communication and is particularly prevalent in insects. However, it has been notoriously difficult to decipher how exactly information related to sexual signaling is encoded in complex chemical profiles. Similarly, our knowledge of the genetic basis of sexual signaling is very limited and usually restricted to a few case studies with comparably simple pheromonal communication mechanisms. The present study jointly addresses these two knowledge gaps by characterizing a single gene simultaneously impacting sexual attractiveness and complex chemical surface profiles in parasitic wasps. Knocking down a fatty acid synthase gene in female wasps dramatically reduces their sexual attractiveness coinciding with a drastic decrease in male courtship and copulation behavior. Concordantly, we found a striking shift of methyl-branching patterns in the female surface pheromonal compounds, which we subsequently demonstrate to be the main cause for the greatly reduced male response. Intriguingly, this suggests a potential coding mechanism for sexual attractiveness mediated by specific methyl-branching patterns, whose genetic underpinnings are not well understood despite their high potential for encoding information. Our study sheds light on how biologically relevant information can be encoded in complex chemical profiles and on the genetic basis of sexual attractiveness.

**Significance Statement:** Unraveling the genetic basis of chemical signaling is one of the most prevalent yet challenging topics in functional genetics and animal communication studies. Here we present the characterization of a biosynthetic gene in parasitoid wasps that simultaneously impacts sexual attractiveness as well as majorly shifts complex surface pheromone compositions. The shifted pattern primarily constitutes up- and down-regulated methyl-branched compounds with very distinct branching positions. Therefore, these findings immediately suggest a potential coding mechanism for sexual attractiveness in complex chemical profiles. This advances our understanding of how genetic information can be translated into biologically relevant chemical information and reveals that sexual attractiveness can have a comparably simple genetic basis.

## Introduction

Tightly coordinated chemical signaling has repeatedly been shown to be of fundamental importance for successful reproduction in a wide range of animal species (1, 2). Particularly insects have exploited this type of signaling as their primary mode of communication (3, 4). Nevertheless, exactly how specific information such as mating status or attractiveness is encoded in the myriad of signaling molecules documented to be involved in sexual communication remains poorly understood (5, 6). Cuticular hydrocarbons (CHCs), major components on the epicuticle of insects, are capable of chemically encoding and conveying a wide variety of biologically relevant information (7, 8). Most prominently, CHCs have been shown to play pivotal roles in sexual communication as the main cues to attract and elicit courtship from conspecific mates (9, 10) to enable discrimination of con- from heterospecific mating partners (11, 12) and to signal receptivity and mating status (13, 14).

Despite such diversified CHC-encoded signals and mediated behaviors, our knowledge on exactly how CHCs encode biologically relevant information has remained surprisingly scarce. This is particularly problematic in studies considering CHC profiles in their entirety as the main signaling entities (13, 15). The exact compounds or their combinations actually encoding the relevant information within CHC profiles remain largely elusive, except for a few case studies mainly involving the dipteran model organism *Drosophila melanogaster*, where single unsaturated CHC compounds appear to be the main mediators in sexual communication (16, 17). In most other cases, chemical information appears to be encoded in a much more complex manner, involving several CHC compounds in different quantitative combinations, with no deeper understanding on the actual coding patterns conveying specific information (9, 18).

In addition to our limited understanding on how CHC profiles encode information, our knowledge of the genetic basis of CHC biosynthesis and its impact on sexual signaling has remained comparably restricted and biased towards the *Drosophila* model system as well (8, 19). In short, the CHC biosynthetic pathway consists of the elongation of fatty-acyl-Coenzyme A units to produce very long-chain fatty acids that are subsequently converted to CHCs (8, 20). An important early switch in CHC biosynthesis is either the incorporation of malonyl-CoA or methyl-malonyl-CoA, eventually leading to the production of straight-chain (*n*-) or methyl-branched (MB-) alkanes, respectively (19, 20, Fig. 1). It has been hypothesized that these processes are mediated by two types of fatty-acyl-synthases (FAS), microsomal for methylmalonyl-CoA and cytosolic for malonyl-CoA (21, 22). To produce olefins (mono- and poly-unsaturated CHCs), desaturases introduce double bonds into straight-chain CHC precursors (8, 19). In *Drosophila*, a couple of genes have been identified that mainly affect the biosynthesis and ratios of unsaturated CHC compounds that function in sexual signaling. For instance, two desaturases (*Desat1, DesatF*) and one elongase (*eloF*) are involved in female diene production and consequently in their functionality as main sex pheromonal compounds (23, 24). Furthermore, in the Australian congeneric species *D. serrata*, the male-specific *D. melanogaster* orthologue *FASN2* has been shown to affect the biosynthesis of three MB-CHCs, among them the additional female mating stimulant 2-Me-C26 (25, 26). Apart from these case studies limited to *Drosophila*, we know very little about the genetic basis linking CHC biosynthesis and sexual signaling in other insects (19).

**Figure 1.**
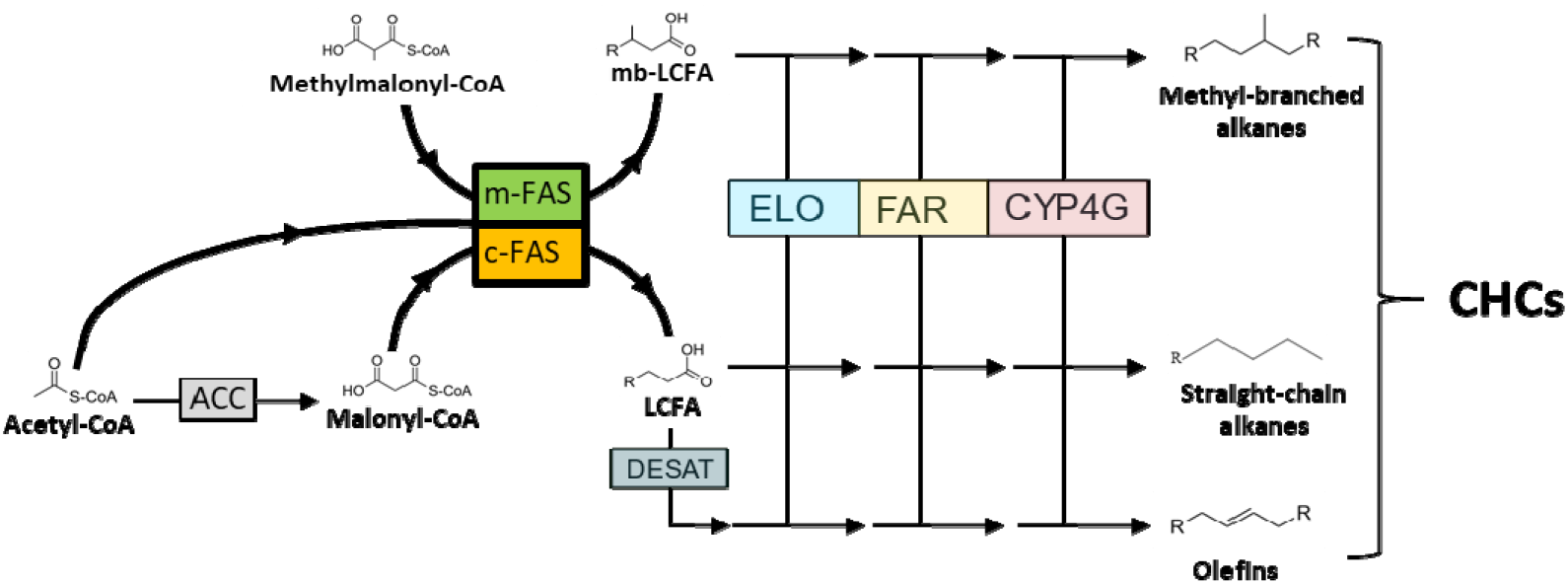
Simplified overview of CHC biosynthesis emphasizing fatty acid synthase (FAS) catalyzed reactions. Initially, acetyl-Coenzyme A (CoA) is converted into malonyl-CoA by the enzyme Acetyl-CoA carboxylase (ACC). Then, further malonyl-CoA subunits are successively incorporated onto the acetyl-CoA primer to form long chain fatty acids (LCFAs) catalyzed by fatty acid synthase enzymes hypothesized to be subcellularly located in the cytosol (c-FAS). For the synthesis of methyl-branched (mb)-CHCs with internal methyl groups, methyl-malonyl-CoA units are incorporated at specific chain locations instead of malonyl-CoA units, catalyzed by m-FAS enzymes whose subcellular location has been hypothesized to be microsomal. The methyl-branched or straight-chain LCFAs are then further processed through a series of biosynthetic conversions catalyzed by the elongase (ELO) enzyme complex, fatty acyl-CoA reductase (FAR) enzymes and cytochrome P450 decarboxylase (CYP4G) enzymes to either methyl-branched or straight-chain, saturated CHCs, respectively. For the biosynthesis of unsaturated CHCs (olefins), desaturase (DESAT) enzymes introduce double bonds into the fatty acyl-CoA chain between elongation steps. For a detailed description of CHC biosynthesis and the involved enzymatic reactions see Blomquist and Ginzel (7) as well as Holze, Schrader and Buellesbach (19).

The parasitoid jewel wasp *Nasonia vitripennis* (Hymenoptera: Pteromalidae) has emerged as a suitable model organism to combine studies on functional genetics as well as chemical communication systems in Hymenoptera (28, 29). Female CHCs serve as sexual cues capable of eliciting male courtship and copulation behavior in *N. vitripennis* (15, 30). The CHC profiles of *N. vitripennis* females exhibit a high complexity, consisting of a mixture of *n*-alkanes, *n*-alkenes and MB-alkanes in various quantities. Specifically the latter fraction, which makes up more than 85 % of the whole profile, displays a rich diversity in methyl-branch numbers, chain lengths and respective relative abundances, hinting at a considerable potential for encoding differential information (18, 30). However, the entire female CHC profile has long been regarded as encoding their sexual attractiveness, with no single compound, compound classes or particular patterns being identifiable as the main conveyers of the sexual signaling function (30, 31). Moreover, despite recent advances in unraveling the genetic architecture of CHC biosynthesis and variation in the *Nasonia* genus (28), the effects of individual genes on CHC profiles and, more importantly, on the encoded sexual signaling function, could not be determined as of yet.

In this study, we characterize the phenotypic effects of a single fatty acid synthase gene (*fas5*) that impacts the variation of several structurally related CHC components concordant with female sexual attractiveness. CHC profiles of *fas5* knockdown females primarily showed significant up- and down-regulations of MB-alkane compounds with correlated branching patterns. At the same time, these knockdown females elicited significantly less courtship and copulation attempts from conspecific males. This constitutes the first hymenopteran gene with a demonstrated function in governing the variation of primarily MB-alkanes as well as sexual attractiveness, hinting at a chemical coding pattern mostly conveyed by this CHC compound class.

## Results

### Knockdown of *fas5* dramatically alters CHC profile composition

RNAi micro-injection into female *Nasonia* pupae at the pupal stage resulted in a striking CHC profile shift in adult females, most prominently displayed in altered MB-alkane patterns (**Fig. 2 A-B, Table S2**). More specifically, the proportions of MB-alkanes with their first methyl branches positioned on the 3^rd^ and 5^th^ C-atom significantly increased in *fas5* RNAi females (13.44% and 47.51%), compared to both WT (11.29% and 16.54%) and GFP RNAi females (10.31% and 16.04%), respectively (**Fig. 2 C-D**). Conversely, MB-alkanes with their first methyl-branches on the 7^th^ C-atom position significantly decreased in *fas5* knockdowns (2.69%) compared to WT (14.16%) and GFP controls (16.31%) (**Fig. 2 E**). Other MB-alkanes with their first methyl-branches mainly positioned on the 9^th^, 11^th^, 13^th^ and 15^th^ also significantly decreased in *fas5* RNAi females (19.02%) as opposed to WT (41.24%) and GFP controls (40.97%) (**Fig. 2 F**). Interestingly, overall MB-alkane amounts as well as total CHC quantity remained stable in *fas5* knockdown females compared to the controls, with no significant overall proportional changes (**Fig. 2 G**). To verify the effect of the knockdown on gene expression levels, we demonstrate a significant reduction in relative *fas5* gene expression in knockdown females compared to both GFP and WT controls **(Fig. 2 H)**. Lastly, *n*-alkene quantities, generally only occurring in negligible quantities in *Nasonia* females (28), also increased significantly in *fas5* RNAi females (3.81%) compared to WT (1.55%) and GFP controls (1.8%) (**Fig. S1 C**). Male CHC profiles were similarly affected by *fas5* knockdowns (**Fig. S2**), however, as there CHCs have not been shown to function as sex pheromones in *N. vitripennis*, those were not investigated further.

**Figure 2.**
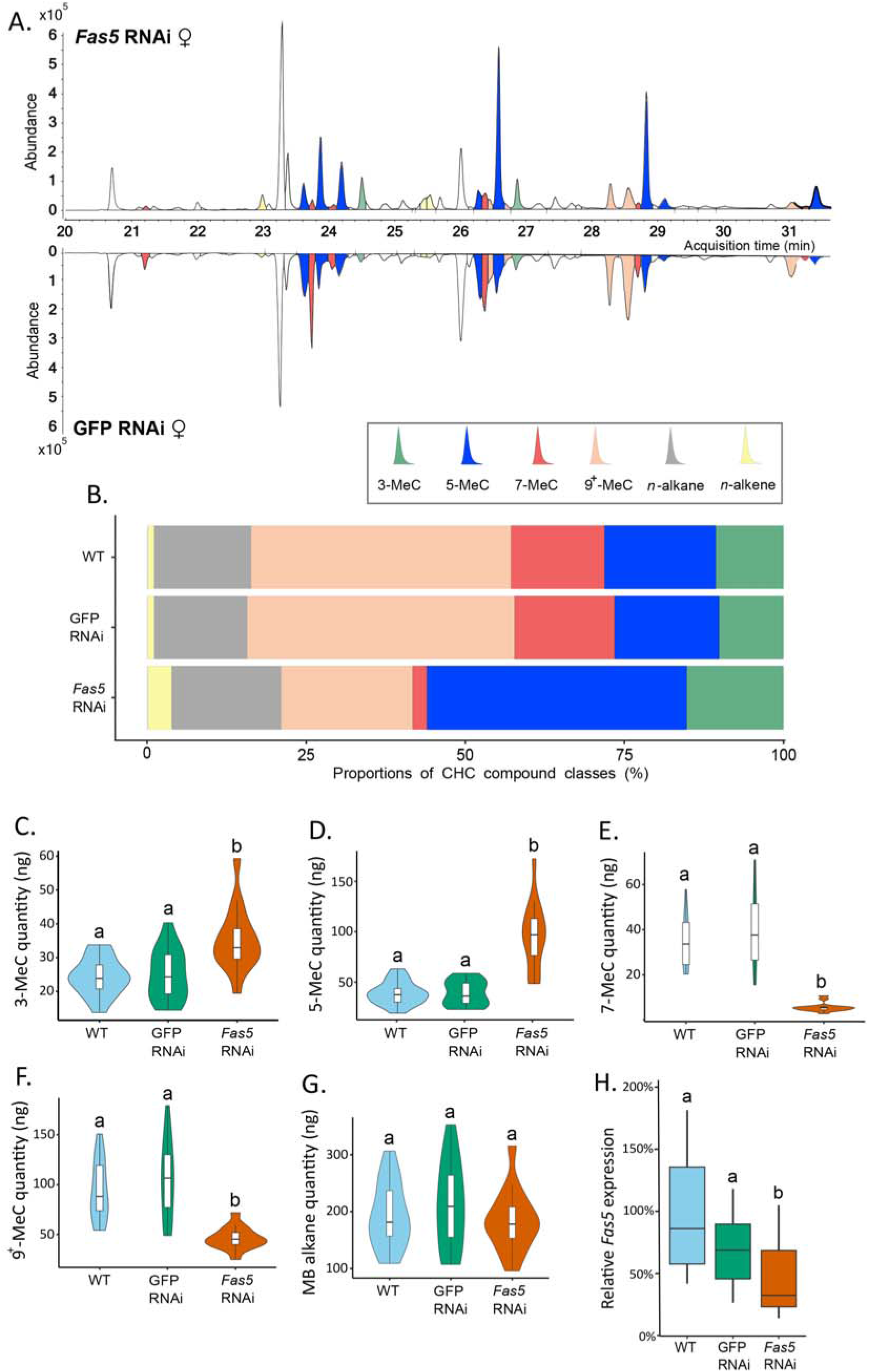
*Fas5* knockdown primarily alters quantities and ratios of methyl-branched CHCs with specific branching patterns in females. **A)** Chromatogram comparison of surface extracts from single female *N. vitripennis* wasps injected with *fas5* dsRNA (top) and GFP dsRNA (bottom). CHC compound peaks with significantly different quantities in *fas5* knockdown vs. GFP control females are indicated in color (compare to Table S2). Different colors are used for methyl-branched (MB) alkanes with their first methyl group at positions 3-, 5-, 7- and 9^+^ (also including positions 11-, 13- and 15-) as well as *n*-alkanes and *n*-alkenes; **B)** Average relative abundances (%) of different CHC compound classes (as indicated in A) compared between wildtype (WT, N=14), control knockdown (GFP, N=15) and *fas5* knockdown (*fas5*, N=15) female wasps; **C)** Average absolute quantities (in ng) of MB alkanes with their first methyl group at the 3^rd^ C-atom position (3-MeC) compared between wildtype (WT), control knockdown (GFP) and *fas5* knockdown (*fas5*) female wasps, indicated by blue, green and orange violin plots, respectively, from here on, sample sizes as in B; **D)** Average absolute quantities of MB alkanes with their first methyl group at the 5^th^ C-atom position (5-MeC), violin plot colors and group designations as in C ; **E)** Average absolute quantities of methyl-branched CHCs with their first methyl group at the 7^th^ C-atom position (7-MeC); **F)** Average absolute quantities of methyl-branched CHCs with their first methyl group at the 9^th^ (as well as 11^th^, 13^th^ and 15^th^, indicated as 9^+^-MeC) C-atom position; **G)** Average absolute quantities of total MB alkane amounts; **H)** Relative expression of *fas5* in WT, GFP and *fas5* RNAi females (N=15 in each treatment), indicated by blue, green and orange boxplots, respectively. Significant differences (p < 0.05) were assessed with Benjamini-Hochberg corrected Mann-Whitney U tests in **A), C)** – **H)** and are indicated by different letters.

### *fas5* knockdown decreases attractiveness of female CHC profiles

*Fas5* RNAi and control (WT and GFP) females elicited antennation from similar proportions of males, while a significantly reduced proportion of males performed courtship and copulation towards *fas5* RNAi females (∼ 50%) compared to controls (∼ 90%) (**Fig. 3 B**). Furthermore, 70 % of the males rejected *fas5* RNAi females at least once, a behavior which was not present at all towards control females (**Fig. 3 C**). To further minimize active female involvement in mate choice and increase male reliance on chemical cues (15, 18), we subsequently offered differentially treated female dummies (*i*.*e*., freeze-killed females) to WT males. Similar proportions of males showed antennation towards freeze killed *fas5* RNAi, GFP RNAi and WT females (**Fig. 3 D**). However, significantly fewer males performed courtship and copulation behavior towards freeze killed *fas5* RNAi females (20%) compared to both controls (> 75%, respectively) (**Fig. 3 D**). To further test whether this dramatic reduction in sexual attractiveness is related to the altered chemical profile, we proceeded to manipulate the CHC profile of female dummies. Female dummies either had their chemical profiles completely removed (*i*.*e*., cleared) or we reconstituted initially cleared WT female dummies with chemical profiles from *fas5* RNAi, GFP RNAi, or WT females, respectively. Likewise, no significant difference was found on the proportion of males that performed antennation towards female dummies of all treatments (**Fig. 3 E**). However, significantly less males performed courtship towards female dummies that were cleared of their chemical profiles (0 %) and *fas5* RNAi re-constituted dummies (16%), compared to WT (50%) and GFP RNAi-reconstituted (75%) female dummies (**Fig. 3 E**). Furthermore, copulation attempts were initiated by less than 5 % of the males towards *fas5* RNAi reconstituted dummies, which is similar to completely cleared dummies (0%). These numbers were in both cases significantly lower compared to control dummies reconstituted with WT and GFP RNAi profiles (∼ 40%, respectively) (**Fig. 3 E**). This demonstrates that the dramatic reduction in female attractiveness in *fas5* knockdown females is predominantly mediated by their altered chemical profiles.

**Figure 3.**
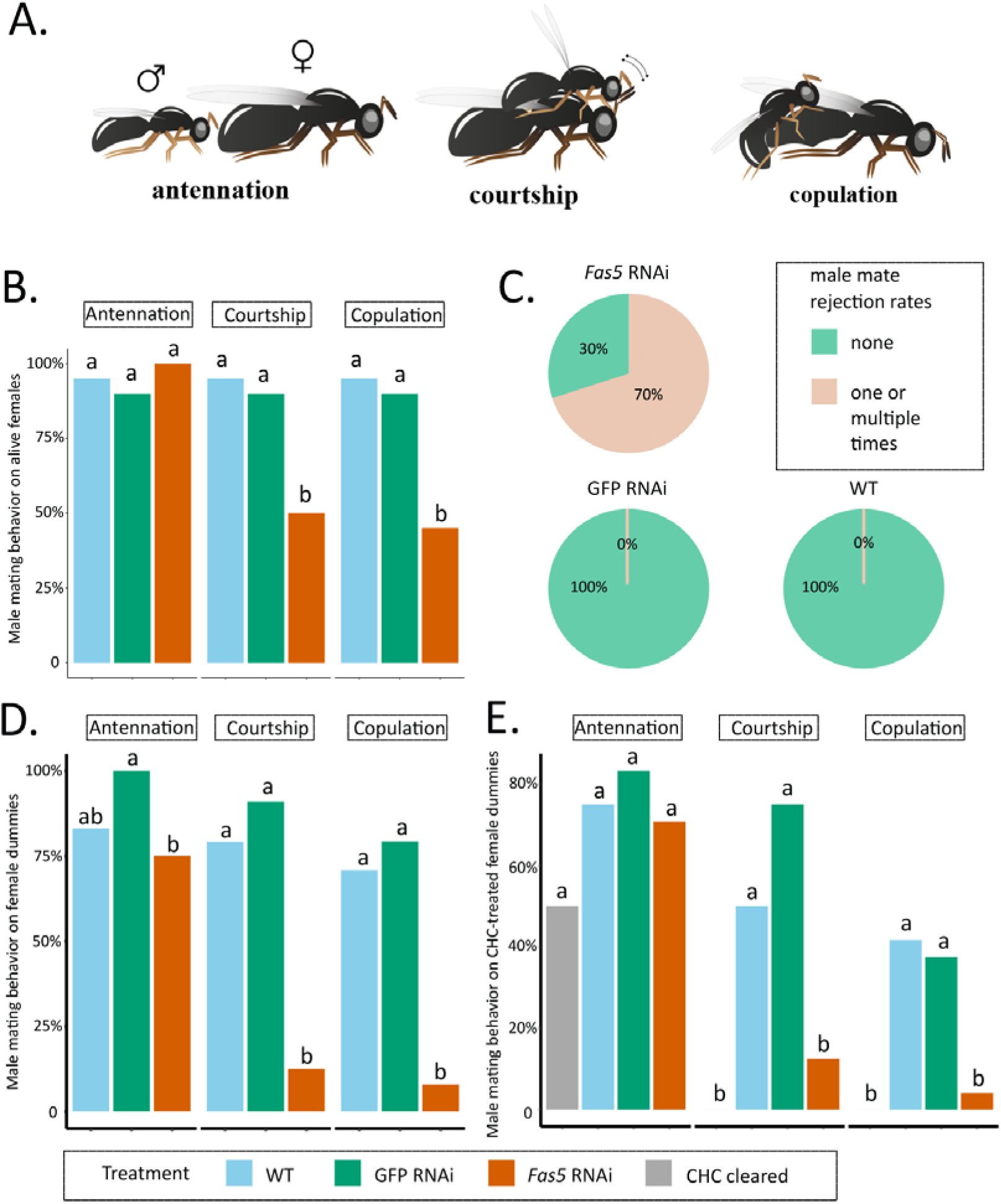
*Fas5* RNAi females elicit less courtship and copulation behaviors from WT males. **A)** Depiction of consecutively displayed mating behavior of *N. vitripennis* males towards females, consisting of initial antennation, courtship (stereotypical head-nods on the female antennae after mounting) and actual copulation (injection of the male aedeagus into the female’s genital opening) (images by Quoc Hung Le). **B)** The proportions of males performing antennation, courtship and copulation towards alive WT, GFP and *fas5* RNAi females, which are marked by blue, green and orange bar plots, N=20 for each treatment; **C)** Male mate rejection rates towards alive WT, GFP and *fas5* RNAi females, separated between no (light green) and one or multiple rejections (light orange); **D)** The proportions of males performing antennation, courtship and copulation towards freeze-killed WT, GFP and *fas5* RNAi females. Bar plot colors and group designations as in B, N=24 for each treatment; **E)** The proportions of males performing antennation, courtship and copulation towards either CHC cleared (in grey) female dummies, or CHC cleared female dummies reconstituted with one female CHC profile equivalent from WT, GFP and *fas5* RNAi females (treatment colors and group designations as in B and D), N=24 for each treatment. Significant differences (p<0.05) were assessed with Benjamini-Hochberg corrected Fisher’s exact tests in **B), D)** - **E)** and are indicated by different letters.

### Female sexual attractiveness is mainly governed by MB-alkanes

To further pinpoint the part of the chemical profile that is responsible for encoding sexual attractiveness, we fractionated the chemical profiles of *N. vitripennis* females and focused on the MB-alkane fraction, which displayed the most conspicuous changes in *fas5* knockdown females (See **Fig. 2 B**). The separation process reduced both the *n*-alkane and *n*-alkene proportions to less than 1% of the whole profile, allowing to focus almost exclusively on the MB-alkane fraction (**Fig. 4 A-B**). Specifically, the MB-alkane fraction of *fas5* RNAi females maintained the dramatically higher proportions of alkanes with 3^rd^ and 5^th^ C-atom methyl-branch positions and lower proportions of alkanes with 7^th^ and 9^th^, 11^th^, 13^th^ and 15^th^ C-atom methyl-branch positions, compared to the respective control females’ MB-alkane fractions (**Fig. 4 C**). We also reconstituted cleared WT female dummies with the separated MB-alkane fractions and offered them to WT males in further behavioral assays. As in our previous behavioral assays with female dummies and reconstituted whole CHC extracts, the MB-alkane fraction from *fas5* RNAi females elicited significantly less courtship and copulation attempts from WT males than the MB-alkane fractions of both WT and GFP RNAi females (**Fig. 4 D**). Overall, these results demonstrate that sexual attractiveness appears to be mainly encoded in the MB-alkane fraction of *N. vitripennis* females.

**Figure 4.**
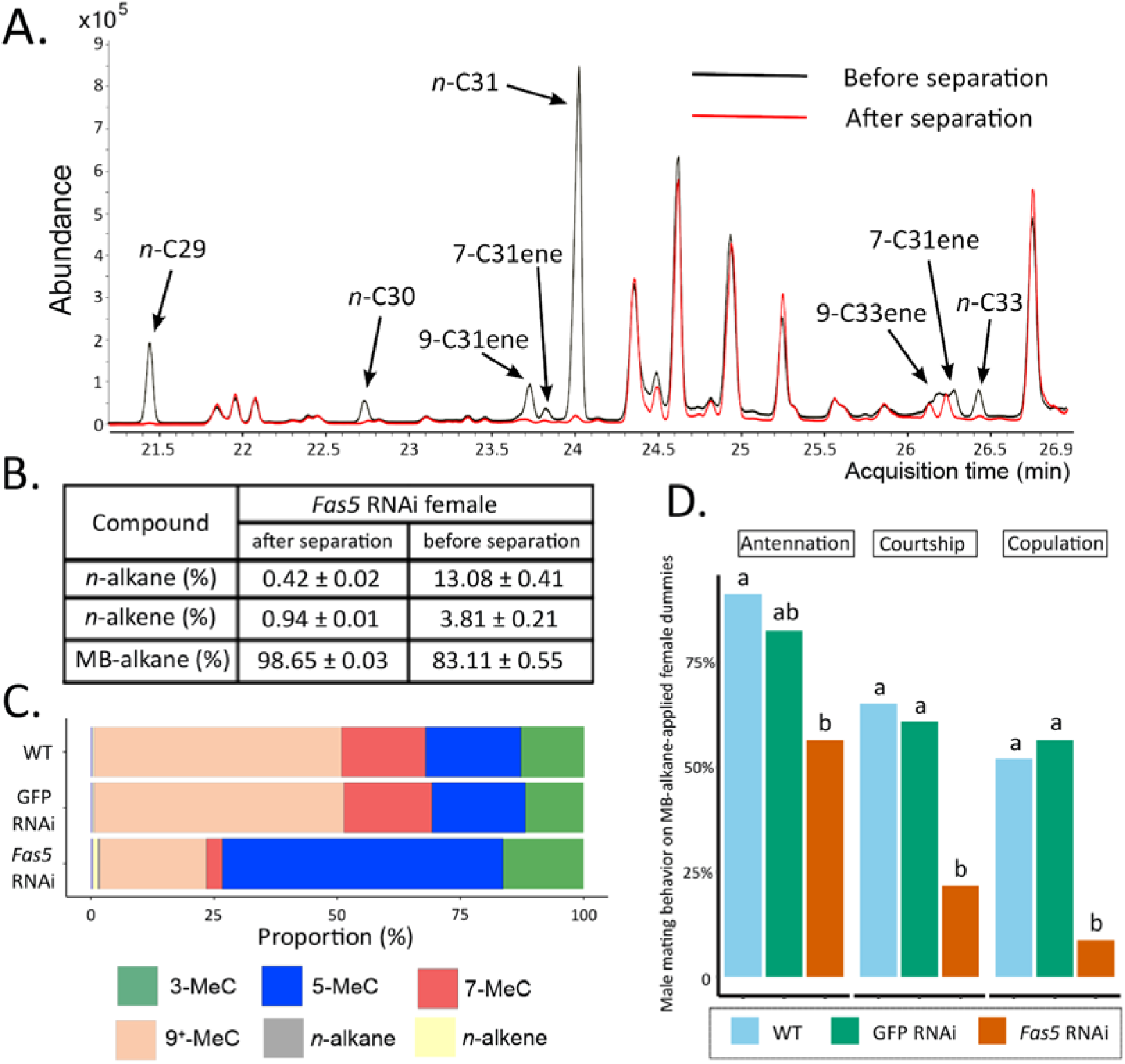
Methyl-branched alkane fraction from *fas5* RNAi females elicits less courtship and copulation from WT males. **A)** Chromatogram comparison of representative *fas5* RNAi female CHC profiles before (in black) and after (in red) physical separation of the methyl-branched (MB) alkane fraction from the other compound classes (*n*-alkanes and *n*-alkenes). Individual *n*-alkane and *n*-alkene compound peaks are indicated by arrows (all other peaks correspond to MB-alkanes). Note that only the part of the *Nasonia* CHC profile where these compounds do occur is shown (compare to Fig. 1 A); **B)** The average relative abundance (%) of overall MB-alkanes, *n*-alkanes and *n*-alkenes in *fas5* RNAi female CHCs before and after separation of the methyl-branched (MB) alkane fraction from the other compound classes. N=3 for each treatment; **C)** Average relative abundances (%) of different compound classes, including MB alkanes with their first methyl group at positions 3-, 5-, 7- and 9^+^ (also including positions 11-, 13- and 15-) as well as *n*-alkanes and *n*-alkenes, compared between wildtype (WT, N=3), control knockdown (GFP, N=3) and *fas5* knockdown (*fas5*, N=3) individuals; **D)** The proportions of males performing antennation, courtship and copulation towards CHC cleared female dummies reconstituted with approximately one female equivalent of MB-alkane fractions derived from WT (in blue, N=23), GFP (in green, N=23) and *fas5* RNAi (in orange, N=24) females. Significant differences (p<0.05) were assessed with Benjamini-Hochberg corrected Fisher’s exact tests in **D)** and are indicated by different letters.

## Discussion

In our study, we shed light on how sexual attractiveness can be encoded by differentially branched cuticular hydrocarbons (CHCs) and unravel its genetic architecture to be based on a single gene. Knocking down the fatty acid synthase gene *fas5* in *Nasonia* wasps leads to a consistent pattern of primarily up- and down-regulated methyl-branched (MB-) alkanes with specific branching patterns. This dramatic shift is accompanied by a significant reduction of courtship and copulation behavior towards knockdown females by conspecific males, which we demonstrate to be mainly determined by the altered MB-alkane fraction. This advances our understanding of how genetic information is translated into chemical information and brings us a step closer in decoding complex chemical profiles.

Most conspicuously, quantities of CHCs with their first methyl groups at the 7^th^ and 9^th^ (and higher) C-atom positions are dramatically down-regulated, whereas the ones with their first methyl groups at the respective 3^rd^ and 5^th^ positions appear mainly up-regulated, in most cases by several orders of magnitude (**Fig. 2** and **Table S2**). Intriguingly, overall CHC quantities do not differ between knockdown and control individuals (**Fig. S1 A**), suggesting a very specific regulatory function for *fas5* in governing these opposing and potentially compensatory branching patterns. Concordantly, this emphasizes the pivotal role of both this particular gene and the wild-type MB-alkane branching patterns in encoding and maintaining the attractiveness of female *N. vitripennis* CHC profiles. Conversely, in the insect model system *Drosophila melanogaster*, CHC-based sexual signaling mechanisms have long been assumed to be comparatively simple, mainly mediated by two doubly unsaturated dienes in females and a mono-unsaturated *n*-alkene in males (16, 17). However, experimental evidence has accumulated that other CHC compounds can also complement sexual signaling in *Drosophila* in various ways (26, 32). When regarding CHC profiles in their entirety as opposed to single compounds, direct causal links between sexual attractiveness and CHC profiles properties have rarely been experimentally demonstrated and have most often defied clear patterns (9, 12). In the genus *Nasonia* and other related parasitoid wasp species, it has so far been assumed that sexual attractiveness is a trait attributed to their entire CHC profile as either present or absent depending on the studied species and also potentially reinforced by other factors such as polar cuticular compounds (18, 33). Our study clearly shows that CHC-mediated sexual attractiveness is primarily conveyed through a relatively complex chemical pattern with a comparatively simple genetic basis.

Interestingly, knockdown of *fas5* also upregulated *n*-alkene quantities (**Fig. S1 C**), which have recently been shown to have a repellent effect on *N. vitripennis* males, preventing them from engaging in homosexual courtship behavior (34). However, our CHC compound class separation greatly reduced the increased proportion of *n*-alkenes in *fas5* knockdown females to levels almost equivalent to those found in wild-type females (*fas5* MB fraction: 0.94 %, WT: 0.91 % in total, compare **Fig. 4 B** to **Table S2**). This renders the contribution of *n*-alkenes to the sharp reduction in sexual attractiveness in *fas5* knockdown females unlikely and strongly suggests that the female sexual signaling function is mainly mediated by MB-alkanes (**Fig. 4**). We argue that this CHC compound class indeed possess the highest potential for encoding a wide variety of chemical information through the myriad of possible positions and numbers of methyl branches. In fact, a couple of studies have already hinted at MB-alkanes as the main carriers for chemical information in insect CHC profiles, providing evidence for the involvement of MB-alkanes in chemical communication processes (35, 36). This might be of particular importance in Hymenoptera, an insect order with CHC profiles largely dominated by MB-alkanes (37, 38) and in which both theoretical considerations as well as empirical evidence have accumulated for the increased complexity and high sophistication of their chemical communication systems (39, 40). Contrary to this, it has been argued that olefins (mainly *n*-alkenes and dienes) have a higher potential for encoding chemical information than MB-alkanes (41). However, this view might have been biased and mainly informed by findings from *Drosophila*, where unsaturated compounds appear to be, in fact, the main mediators of sexual communication (16, 42). In direct comparison, MB-alkanes constitute the dominant fraction in *Nasonia* CHC profiles (> 85 %) (28) as opposed to *Drosophila* profiles (16-24 %) (43). Since the split between Hymenoptera and other holometabolous insects including Diptera has been estimated to have occurred 327 mya (44), fundamental shifts in basic properties of both surface profile compositions as well as chemical signaling functionalities might be expected. Therefore, the promising role of MB-alkanes in conveying chemical information should be investigated more prominently in future studies. Through a wider evolutionary lens, it will be interesting to investigate how the present findings compare to other communication systems with a different chemical basis, for instance in moths, where species-specific sexual attractiveness is largely mediated by ratios of long-range pheromonal components such as alcohols, aldehydes, and acetate esters (2, 6).

Concerning gene orthology, *fas5* has been annotated as a homolog to *FASN3* in *D. melanogaster* (28, 45). Interestingly, knockdowns of *FASN3* alone do not induce any compound changes in *D. melanogaster* CHC profiles but increase the flies’ sensitivity to desiccation (25). A *FASN3* ortholog expressed in the kissing bug *Rhodinius prolixus* also contributes to desiccation resistance, but simultaneously down-regulates MB-CHCs while up-regulating straight-chain CHCs (46). Similarly, in the migratory locust *Locusta migratoria*, silencing two *FAS* genes decreased insect survival under desiccation stress while altering the amounts of both MB- and straight-chain CHCs (47). In contrast to these studies, we were not able confirm any functionality in desiccation resistance for *fas5* in both *N. vitripennis* males and females (**Fig. S3**), nor did the previous studies report any impact on CHC-based chemical signaling. There are two additional *FAS* genes characterized in *D. melanogaster*: *FASN1*, which is responsible for overall CHC production with no specific effects on any particular compound classes (25) and *FASN2*, which exclusively regulates MB-alkane production in males (26). Although not a direct homolog, the main impact on MB-alkane variations of *FASN2* in *D. melanogaster* is comparable to that of *fas5* in *N. vitripennis*, with the exception that we were able to document this effect for both sexes (compare **Fig. 2** and **Fig. S2**). Such diversified functionalities of the *FAS* genes characterized so far indicate their high versatility as early mediators in the CHC biosynthesis pathway (**Fig. 1**). *FAS* genes have been implied as instrumental for generating the huge diversity of different CHC profiles across insects, with high evolutionary turnover rates and differences in the specific functional recruitments of *FAS* gene family members (46, 48). However, *FAS* genes are far from restricted to impact CHC biosynthesis alone and have been documented to be involved in a wide variety of other physiological processes, ranging from lipogenesis to diapause induction (45, 49). Therefore, specifically predicting functionalities of *FAS* genes and unambiguously associating them with CHC biosynthesis and variation has been notoriously difficult (19). Our study suggests a very specific affinity of *fas5* for particular methyl branching patterns, potentially originating in an enzymatic preference for methyl-malonyl-CoA predecessors with pre-existing branching patterns (8, 20). To clarify this, future studies should determine whether the *fas5* gene product constitutes a soluble cytosolic or membrane-bound microsomal FAS enzyme, the latter of which has been postulated to be specific for the incorporation of methyl-malonyl CoA subunits (21, 22, Fig. 1).

In conclusion, our study demonstrates the considerable impact of a single fatty acid synthase gene on female sexual attractiveness in a parasitoid wasp, thereby immediately suggesting how this trait can be encoded through specific ratios of MB-alkanes. To the best of our knowledge, *fas5* is the first identified hymenopteran gene with a specific effect on MB-alkane ratios that simultaneously impacts sexual attractiveness. Transcending the demonstrated impact on sexual signaling and elicited mating behavior, the present findings also substantially advance our general knowledge on the so far little investigated genetic underpinnings of MB-alkane variation and production. This particular compound class dominates the surface profiles of many insects, most notably in the ecologically and economically prevalent order Hymenoptera, and harbors a considerable potential for encoding chemical information, inviting a stronger emphasis on these compounds in future studies on chemical signaling and its behavioral impact.

## Materials and Methods

### *Nasonia* strain maintenance and preparation

The standard laboratory strain AsymCX of *Nasonia vitripennis*, originally collected in Leiden, the Netherlands, was used for all experiments. The wasps were reared under 25°C, in 55% relative humidity and a light:dark cycle of 16:8h, leading to a life cycle of ∼14 days. Pupae of *Calliphora vomitoria* (Diptera: Calliphoridae) were used as hosts.

### RNAi gene knockdown

For the knockdown of *fas5*, dsRNA was synthesized following the manual of MEGA Script T7 kit (Invitrogen, Carlsbad, CA, USA), using the primer pairs listed in **Table S1**. The Quick-RNA Tissue/Insect Kit (Zymo Research, Freiburg, Germany) was used to purify the resulting dsRNA product. GFP (Green Fluorescent Protein) dsRNA, which has no known targets in the *Nasonia* genome (50), was used as a control. GFP dsRNA was synthesized from the vector pOPINEneo-3C-GFP, which was kindly donated by Ray Owens (Addgene plasmid #53534; http://n2t.net/addgene:53534; RRID:Addgene_53534). Microinjections were performed with 4-5 µg/µl (diluted in nuclease free water, Zymo Research) *fas5* and GFP dsRNA on a Femtojet microinjector (Eppendorf, Hamburg, Germany) following the protocol published by Lynch et al (51). *N. vitripennis* yellow pupae (7 to 8 days old after egg deposition) were gently fixed on a cover slide using double-sided tape (Deli, Zhejiang, China), with their abdomens facing up. DsRNA mixed with 10% red food dye (V2 FOODS, Niedersachsen, Germany) was injected into the abdomens of the pupae using a thin needle, which was produced in a PC-10 puller (Narishigne Group, Tokyo, Japan) by heating a glass capillary (100 mm length x 85 µm inner diameter, Hilgenberg, Malsfeld, Germany) to 100°C, and subsequently breaking the stretched capplilary in two parts in the narrow middle section at 67 °C. Individual injections were performed until the red dye has spread evenly within the abdomen of each pupa. The injected pupae were then stored inside a petri dish with a piece of wet tissue at the bottom to ensure saturation with sufficient humidity for the pupae to mature and eclose. After the pupae elosed as adults, they were collected at an age of 0-24 h and snap-frozen with liquid nitrogen, after which they were stored at -80 °C for further experiments.

### RNAi efficiency analysis

RNAi knockdown efficiency was determined by quantitative PCR (qPCR), assessing *fas5* gene expression levels between knockdown and control individuals. RNA from each individual wasp after chemical extraction (see below) was obtained using the Quick-RNA Tissue/Insect Kit (Zymo Research, Freiburg, Germany), and reversely transcribed into complemetary DNA (cDNA) utilizing the cDNA Synthesis Kit (CD BioSciences, New York, USA). As controls for the qPCR procedure, we used *N. vitripennis elongation factor 1*α (*NvEF-1*_α_) as a housekeeping gene, as described by Wang, *et al*. (34). The qPCR was performed in a Lightcycler480 qPCR machine (Roche, Basel, Switzerland), with a pre-incubation of 95°C for 3 minutes, 40 amplification cycles of 15 seconds at 95°C and 60 seconds of 60°C, as well as a final standard dissociation curve step to check the specificity of the amplification.

### Chemical analysis

Chemical extractions of single wasps were performed by immersing them in 50 µl HPLC-grade *n*-hexane (Merck, KGaA, Darmstadt, Germany) in 2 ml glass vials (Agilent Technologies, Waldbronn, Germany) on an orbital shaker (IKA KS 130 Basic, Staufen, Germany) for 10 minutes. Extracts were subsequently evaporated under a constant stream of gaseous carbon dioxide and then resuspended in 10 μl of a hexane solution containing 7,5 ng/μl dodecane (C12) as an internal standard. Following this, 3 μl of the resuspended extract was injected in splitless mode with an automatic liquid sampler (ALS) (PAL RSI 120, CTC Analytics AG, Switzerland) into a gas-chromatograph (GC: 7890B) simultaneously coupled to a flame ionization detector (FID: G3440B) and a tandem mass spectrometer (MS/MS: 7010B, all provided by Agilent Technologies, Waldbronn, Germany). The system was equipped with a fused silica column (DB-5MS ultra inert; 30 m x 250 μm x 0.25 μm; Agilent J&W GC columns, Santa Clara, CA, USA) with helium used as a carrier gas under a constant flow of 1.8 ml/min. The FID had a temperature of 300 °C and used nitrogen with a 20 mL/min flow rate as make-up gas, and hydrogen with a 30 mL/min flow rate as fuel gas. The column was split at an auxiliary electronic pressure control (Aux EPC) module into an additional deactivated fused silica column piece (0.9 m x 250 μm x 0.25 μm) with a flow rate of 0.8 mL/min leading into the FID detector, and another deactivated fused silica column piece (1.33 m x 250 μm x 0.25 μm) at a flow rate of 1.33 mL/min leading into the mass spectrometer. The column temperature program started at 60 °C and was held for 1 min, increasing 40 °C per minute up to 200 °C and then increasing 5 °C per minute to the final temperature of 320 °C, held for 5 min.

CHC peak detection, integration, quantification and identification were all carried out with Quantitative Analysis MassHunter Workstation Software (Version B.09.00 / Build 9.0.647.0, Agilent Technologies, Santa Clara, California, USA). CHCs were identified according to their retention indices, diagnostic ions, and mass spectra as provided by the total ion count (TIC) chromatograms, whereas their quantifications were achieved by the simultaneously obtained FID chromatograms, allowing for the best-suited method for hydrocarbon quantification (Agilent Technologies, Waldbronn, Germany, pers. comm.) while simultaneously retaining the capability to reliably identify each compound. Absolute CHC quantities (in ng) were obtained by calibrating each compound according to a dilution series based on the closest eluting *n*-alkane from a C21-40 standard series (Merck, KGaA, Darmstadt, Germany) at 0.5, 1, 2, 5, 10, 20, 40 ng/µl, respectively.

### CHC compound class separations

Physical separation of the methyl-branched (MB-) alkane fraction from the other compound classes in *N. vitripennis* CHC profiles was performed according to an adapted protocol from Würf, Pokorny, Wittbrodt, Millar and Ruther (9) and Bello, McElfresh and Millar (52). CHC profiles of approximately 700 females were extracted in 9 mL HPLC-grade *n*-hexane (Merck, KGaA, Darmstadt, Germany) which was subsequently evaporated under a stream of gaseous carbon dioxide. The dried extract was re-suspended in 10 mL isooctane (99%, Sigma-Aldrich, Taufkirchen, Germany), and stirred overnight with a magnetic stirrer (Model: C-MAG HS4, IKA, Germany) after adding 2 g activated (*i*.*e*., baked at 300°C for 2h) molecular sieves (5Å, 45-60 mesh size, Merck, KGaA, Darmstadt, Germany). The molecular sieves were filtered out by loading the extract into a glass funnel (50 mm inner diameter) with glass wool (Merck, KGaA, Darmstadt, Germany) and 0.2 g silica gel (High purity grade, pore size 60Å, 230-400 mesh particle size, Merck, KGaA, Darmstadt, Germany) containing 10 % pulverized AgNO_3_ (99.7%, Merck KGaA, Darmstadt, Germany). This procedure is devised to effectively filtering out *n*-alkanes and olefins, retaining only the MB-alkane fraction in the remaining extract (9, 52). The isooctane in the extract was condensed to 2 mL under a stream of gaseous carbon dioxide, from which we sampled 50 µl to estimate the quantity of the overall extract. Based on the quantity of overall MB-alkanes and single female CHC profiles, the MB-alkanes were reconstituted in hexane to a final concentration of approximately one female equivalent per 5 µl, which was saved for further behavioral assays.

### Behavioral assays

Mating behavior assays were carried out to test whether female sexual attractiveness decreased after knockdown of *fas5*. First, virgin females of three treatments (*fas5* RNAi, GFP RNAi and WT) were offered to 0-48 h old virgin WT *N. vitripennis* (AsymCX) males, following the protocol described in Wang, *et al*. (34). A female was transferred into a transparent plastic vial (76mm height, 10mm diameter) that contained a male. The assay was started as soon as the male was introduced and observed for 5 mins. The males’ behavior towards the females was then scored based on presence or absence of three consecutive behavioral displays: antennation (physical contact of male antennae with the female’s body surface), courtship (series of stereotypic headnods and antennal sweeps after mounting the female) and actual copulation attempts, which have been established as indicators of male mate acceptance and female sexual attractiveness (15, 53, Fig. 3 A). If the male did not initiate any further courtship or copulation behaviors after antennation, this was scored as a male mate rejection as described in Buellesbach, Greim and Schmitt (53).

To increase the focus on the male mate choice behavior in relation to female chemical cues, further behavioral assays were carried out with freeze-killed females (*i*.*e*., dummies) offered to 0-48 h old virgin WT males. In the first set of these experiments, male behavior was recorded and compared on female dummies of the three previously mentioned treatments (*fas5* RNAi, GFP RNAi and WT). In the second set, female dummies were manipulated by either soaking them individually in 50 µl hexane for 3 h, effectively removing their CHC profiles (15, 18) or reconstituting soaked WT female dummies with one female CHC profile equivalent (resuspended in 5 µl hexane as described above for chemical analysis) from *Nv_fas5* RNAi, GFP RNAi and WT females. In the third set, approximately one female equivalent of the separated methyl-branched alkane fraction, prepared directly after the CHC separation process (see above), was reconstituted to CHC cleared female dummies, and offered to the WT males.

All behavioral assays with female dummies were performed in a mating chamber which consisted of two identical aluminium plates (53 × 41 × 5 mm). Each plate contained 12 holes (6 mm diameter) that served as observation sites. In preparation for the recording of the behavioral assays, single female dummies were placed into each hole in one plate, while single WT males were placed into each hole in the opposite plate and immediately covered with glass slides (Diagonal GmbH &Co. KG, Münster, Germany). The behavioral assays were initiated by quickly adjoining the two plates. Recordings were conducted with a Canon camera (EOS 70D, Tokyo, Japan) for 5 mins. All behavioral assays were performed in an eclosed wooden box with constant illumination (100Lm, LED light L0601, IKEA Dioder, Munich, Germany).

### Desiccation assays

Since CHC profiles also play a pivotal role in desiccation prevention (8), we further performed desiccation assays to explore the impact of *fas5* knockdowns on the wasps’ ability to survive under different degrees of desiccation stress. High desiccation stress conditions were implemented by placing 0.6 g desiccant (DRIERITE, Merck, KGaA, Darmstadt, Germany) into transparent plastic vials (76mm height, 10mm diameter), which were airtightened with rubber plugs, resulting in low (9%) relative humidity after 24 h. Inside each vial, a piece of cotton and a stainless-steel grid was placed in the middle to separate the desiccant from the remaining space of the vial (∼ 35 mm height), which served as an observation site where individiual wasps were placed for the duration of the desiccation assay. Control vials with moderate desiccation stress (∼ 55% relative humidity) were prepared similarly but without adding the desiccant. the relative humidity in the test tubes of different humidity treatments was monitored using a humidity-temperature probe (Feuchtemesssystem HYTELOG-USB, B+B Thermo-Technik GmbH, Donaueschingen, Germany) with a measurement accuracy of ± 2% relative humidity at 23°C. Newly eclosed (0-24 h old) male and female wasps were collected, sorted in groups of 10, and fed with honey water (Bluetenhonig, dm-drogerie markt GmbH & Co. KG, Karlsruhe, Germany) for 9 hours. Afterwards, each group of wasps was randomly assigned to the previously prepared vials with either high or moderate desiccation stress, and recordings were performed with a looped VLC media player (VideoLAN, Paris, France) script, initiating a 2 min recording (Logitech C920 HD PRO webcam, Logitech GmbH, München, Germany) every 2 hours, until the last wasp fell down on the grid and stop moving. The numbers of alive wasps in each vial was assessed, based on which survival curves were built and further compared among treatments.

## Acknowledgments

We would like to thank Marek Golian, Stella Schummer, Anastasia Dzemiantsei and Lisa Sewald for assisting the experiments. Besides, we appreciate Yunsheng Zhu’s help in data analysis and Yidong Wang’s insights in figure construction. Furthermore, we thank Sabine Nooten and Erik T. Frank for helpful suggestions on the first draft of this manuscript.

