## Supplemental figures and table for "Decoding the genetic and chemical basis of sexual attractiveness in parasitic wasps"

*Jan Buellesbach.

**This PDF file includes:**

Figures S1 to S3

Table S1 to S2

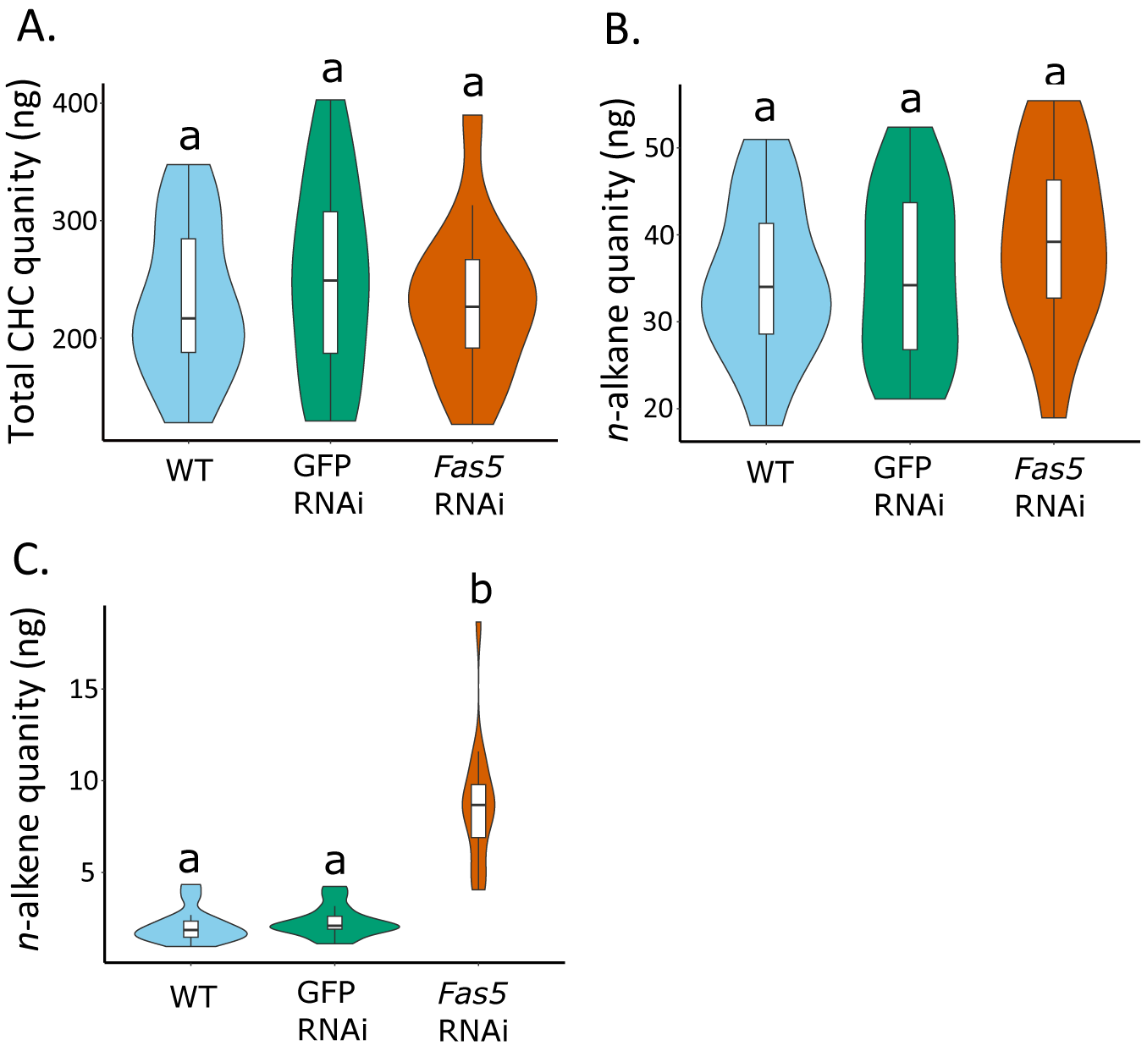

Fig. S1. *Fas5* knockdown does not change total CHC and *n*-alkane quantities but increases *n*-alkene quantities in females. A) Average absolute quantities (ng) of total extracted CHC amounts, compared between wildtype (WT, N=15), control knockdown (GFP, N=15) and *fas5* knockdown (*fas5*, N=14) *N. vitripennis* females, indicated by blue, green and orange violin plots, respectively; B) Average absolute quantities (ng) of the total amount of *n*-alkanes from female CHC extracts, violin plot colors and group designations as in A; C) Average absolute quantities (ng) of the total amount of *n*-alkenes from female CHC extracts, violin plot colors and group designations as in A. Significant differences (p<0.05) were assessed with Benjamini-Hochberg corrected Fisher’s exact tests and are indicated by different letters.

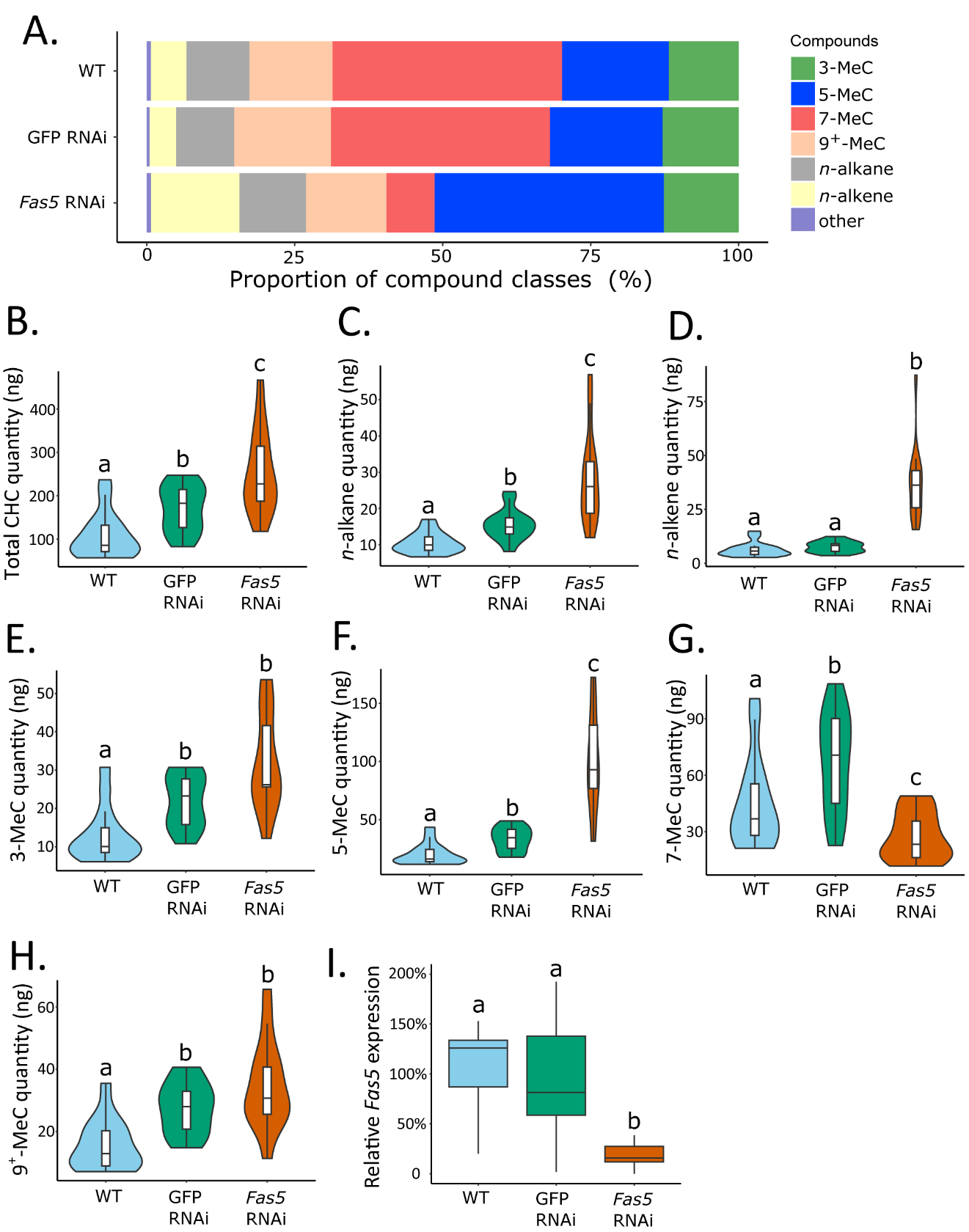

Fig. S2. *Fas5* knockdown primarily alters the ratios of methyl-branched CHCs with specific branching patterns in males. A) Average relative abundances (%) of different CHC compound classes compared between wildtype (WT, N=15), control knockdown (GFP, N=15) and *fas5* knockdown (*fas5*, N=14) male *N. vitripennis* wasps. Different colors are used for methyl-branched (MB) alkanes with their first methyl group at positions 3-, 5-, 7- and 9^+^ (also including positions 11-, 13- and 15-) as well as *n*-alkanes and *n*-alkenes (compare to Fig. 1) B) Average absolute quantities (ng) of total CHC amounts, compared between wildtype (WT), control knockdown (GFP) and *fas5* knockdown (*fas5*) male wasps, indicated by blue, green and orange violin plots, respectively, from here on, sample sizes as in A. C) Average absolute quantities (ng) of total *n*-alkane amounts in male wasps, violin plot colors and group designations as in B D) Average absolute quantities (ng) of total *n*-alkene amounts in male wasps E) Average absolute quantities of MB alkanes with their first methyl group at the 3^rd^ C-atom position (3-MeC); F) Average absolute quantities of MB alkanes with their first methyl group at the 5^th^ C-atom position (5-MeC); G) Average absolute quantities of methyl-branched CHCs with their first methyl group at the 7^th^ C-atom position (7-MeC) H) Average absolute quantities of methyl-branched CHCs with their first methyl group at the 9^th^ (as well as 11^th^, 13^th^ and 15^th^, indicated as 9^+^-MeC) C-atom position; I) Relative expression of *fas5* in WT, GFP and *fas5* RNAi males (N=15 in each treatment), indicated by blue, green and orange boxplots, respectively. Significant differences (p<0.05) were assessed with Benjamini-Hochberg corrected Mann-Whitney U tests and are indicated by different letters.

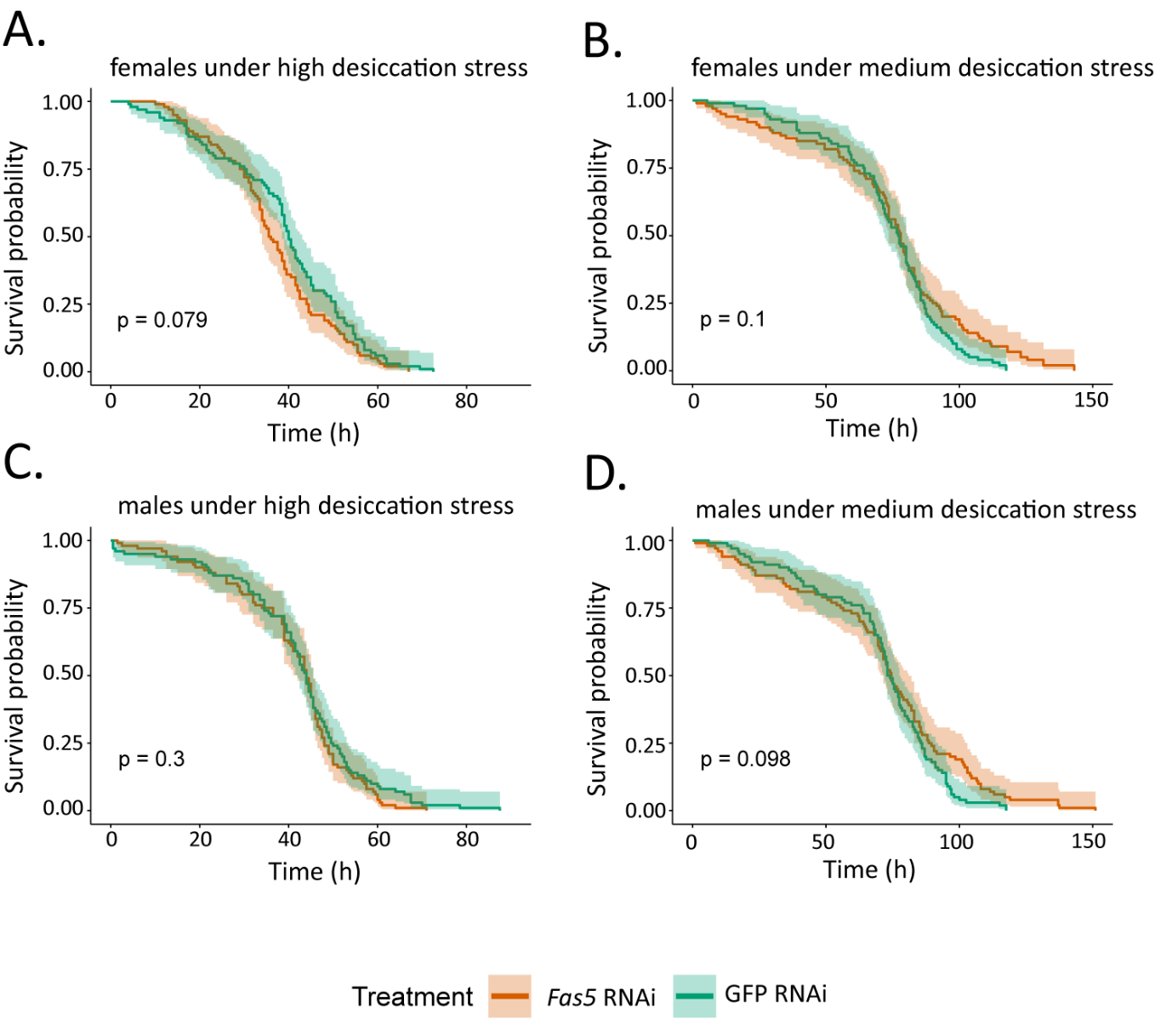

Fig. S3. *Fas5* knockdown does not change survival times of male and female wasps under desiccation stress. A) Comparison of survival probabilities along the observation time under high desiccation stress between control knockdown (GFP RNAi) and *fas5* knockdown (*fas5* RNAi) females; B) Comparison of survival probabilities along the observation time under medium desiccation stress between GFP RNAi and *fas5* RNAi females; C) Comparison of survival probabilities along the observation time under high desiccation stress between GFP RNAi and *fas5* RNAi males; D) Comparison of survival probabilities along the observation time under medium desiccation stress between GFP RNAi and *fas5* RNAi males. N=10 for each treatment. The high desiccation stress treatment was achieved with approximately 9% relative humidity, and the medium desiccation stress treatment with approximately 55% relative humidity as assessed by humidity-temperature probes. Survival probability was assessed with a Kaplan Meier Analysis, and the colored area along the survival curve represents the 95% confident interval.

Table S1. List of primers used in the present study. All primers were designed using the Primer-BLAST tool from the National Center for Biotechnology Information (NCBI). Indicated are the respective primer names, their sequences, and their usage in the experimental protocol.

| **Primer name** | **Sequence** | **Usage** |
| --- | --- | --- |
| *fas5*_dsRNA_F | GATGCAAAGACCAACAAAGCC | dsRNA synthesis for targeting *fas5* |
| *fas5*_dsRNA_R | GCAAAGATTTCGCGATTCCTG |  |
| *fas5*_dsRNA_T7_F | TAATACGACTCACTATAGGGGATGCAAAGACCAACAAAGCC |  |
| *fas5*_dsRNA_T7_R | TAATACGACTCACTATAGGGGCAAAGATTTCGCGATTCCTG |  |
| GFP_dsRNA_F | GTGACCACCTTGACCTACGG | dsRNA synthesis for targeting GFP |
| GFP_dsRNA_R | TCTCGTTGGGGTCTTTGCTC |  |
| GFP_dsRNA_T7_F | TAATACGACTCACTATAGGGGTGACCACCTTGACCTACGG |  |
| GFP_ dsRNA_T7_R | TAATACGACTCACTATAGGGTCTCGTTGGGGTCTTTGCTC |  |
| *fas5*_qPCR_F | CTATGTTTGATGATATCAAGGCTGA | quantitative (q) PCR for *fas5* expression |
| *fas5*_qPCR_R | CATATGTATACTGGTCCCCGTAAG |  |
| *Elf1a*_qPCR_F | GAGCCATCCACCAAAATGCC | quantitative (q) PCR for *Elf1a* expression (*N. vitripennis* elongation factor 1α, housekeeping gene) |
| *Elf1a*_qPCR_R | CTTGTAGACGTCCTGGAGGG |  |

**Table S2.** **Comparison of absolute quantities and relative abundances of single CHC compounds between differentially treated female wasps.** Indicated are retention indices (RI), CHC compound identifications or possible configurations in case of ambiguities, their mean absolute (ng) amounts with their respective absolute standard deviations (sd) as well as their respective relative amounts (in %) compared between wildtype (WT, N=14), control knockdown (GFP RNAi, N=15) and *fas5* knockdown (*fas5* RNAi, N=15) female wasps. Significant effects in *fas5* knockdown (KD) females are indicated by up- (white) and downwards (black) arrows, corresponding to either up- or down-regulation of the absolute compound quantities, respectively. Where compound identifications were ambiguous due to multiple possible methyl branch positions which could be interpreted based on the detected ion pairs, all possible compound configurations are given.

| RI | Compound IDs / configurations | WT female | | GFP RNAi female | | *Fas5* RNAi female | |  |
| --- | --- | --- | --- | --- | --- | --- | --- | --- |
|  |  | mean±sd (ng) | % | mean±sd (ng) | % | mean±sd (ng) | % | KD effect |
| 2900 | n-C29 | 11.11 ± 2.4 | 4,82 | 9.84 ± 2.46 | 3,95 | 11.75 ± 2.34 | 5,06 |  |
| 2939 | 11-MeC29 | 0.94 ± 0.35 | 0,41 | 1.1 ± 0.37 | 0,44 | 1.34 ± 0.81 | 0,58 |  |
| **2947** | **7-MeC29** | **4.12 ± 1.4** | **1,79** | **5.26 ± 2.24** | **2,11** | **1.42 ± 0.79** | **0,61** |  |
| 2956 | 5-MeC29 | 0.7 ± 0.22 | 0,30 | 0.8 ± 0.27 | 0,32 | 1.21 ± 0.66 | 0,52 |  |
| 2977 | 3-MeC29 | 0.29 ± 0.13 | 0,12 | 0.32 ± 0.1 | 0,13 | 0.53 ± 0.29 | 0,23 |  |
| 2982 | 5,17-DiMeC29 | 0.47 ± 0.17 | 0,20 | 0.56 ± 0.2 | 0,22 | 0.7 ± 0.32 | 0,30 |  |
| 3000 | n-C30 | 1.24 ± 0.41 | 0,54 | 1.28 ± 0.44 | 0,52 | 1.39 ± 0.45 | 0,60 |  |
| 3015 | 3,7-; 3,11-DiMeC29 | 0.32 ± 0.07 | 0,14 | 0.37 ± 0.08 | 0,15 | 0.41 ± 0.11 | 0,18 |  |
| 3036 | 3,7,11-; 3,7,13-; 3,7,15-; 3,7,17-TriMeC29 | 0.23 ± 0.1 | 0,10 | 0.31 ± 0.14 | 0,13 | 0.33 ± 0.16 | 0,14 |  |
| **3044** | **7-MeC30** | **0.66 ± 0.37** | **0,28** | **0.91 ± 0.48** | **0,37** | **0.03 ± 0.02** | **0,01** |  |
| **3054** | **5-MeC30** | **0.04 ± 0.02** | **0,02** | 0.05 ± 0.02 | 0,02 | **0.11 ± 0.06** | **0,05** |  |
| **3061** | **4-MeC30** | **0.05 ± 0.02** | **0,02** | **0.05 ± 0.01** | **0,02** | **0.18 ± 0.05** | **0,08** |  |
| **3087** | **9-C31ene** | **0.62 ± 0.34** | **0,27** | **0.83 ± 0.47** | **0,33** | **2.77 ± 1.44** | **1,19** |  |
| **3096** | **7-C31ene** | **0.41 ± 0.15** | **0,18** | **0.39 ± 0.16** | **0,16** | **1.12 ± 0.38** | **0,48** |  |
| 3100 | n-C31 | 21.35 ± 6.47 | 9,26 | 22.93 ± 7.17 | 9,20 | 25 ± 8.76 | 10,77 |  |
| **3134** | **9-; 11-; 13-; 15-MeC31** | **11.54 ± 4.65** | **5,01** | **14.52 ± 6.83** | **5,83** | **6.88 ± 2.97** | **2,97** |  |
| **3140** | **7-MeC31** | **15 ± 6.4** | **6,51** | **19.23 ± 8.86** | **7,72** | **1.38 ± 0.61** | **0,59** |  |
| **3150** | **5-MeC31** | **4.88 ± 1.19** | **2,12** | **4.9 ± 1.53** | **1,97** | **11.52 ± 5.01** | **4,96** |  |
| **3157** | **11,15-; 11,17-; 11,19-; 13,15-; 13,17-; 13,19-DiMeC31** | **0.42 ± 0.34** | **0,18** | **0.4 ± 0.24** | **0,16** | **0.7 ± 2.43** | **0,30** |  |
| **3173** | **7,11-DiMeC31** | **2.2 ± 0.78** | **0,96** | **2.29 ± 1.06** | **0,92** | **0.51 ± 0.2** | **0,22** |  |
| **3178** | **3-MeC31** | **6.05 ± 1.45** | **2,62** | **5.82 ± 2.22** | **2,34** | **10.21 ± 3.75** | **4,40** |  |
| 3187 | 5,15-; 5,17-; 5,19-DiMeC31 | 0.27 ± 0.1 | 0,12 | 0.26 ± 0.1 | 0,11 | 0.18 ± 0.08 | 0,08 |  |
| **3211** | **3,11-; 3,13-; 3,15-DiMeC31** | **1.27 ± 0.37** | **0,55** | **1.42 ± 0.36** | **0,57** | **6.14 ± 2.1** | **2,64** |  |
| 3217 | 3,7-; 3,9-DiMeC31 | 0.73 ± 0.2 | 0,32 | 0.66 ± 0.18 | 0,27 | 0.93 ± 0.22 | 0,40 |  |
| 3236 | 3,7,9-; 3,7,11-; 3,7,15-TriMeC31 | 2.03 ± 0.64 | 0,88 | 2.35 ± 0.92 | 0,94 | 2.12 ± 0.55 | 0,91 |  |
| **3253** | **6-MeC32** | **0.21 ± 0.08** | **0,09** | **0.21 ± 0.1** | **0,08** | **0.04 ± 0.02** | **0,02** |  |
| 3263 | 3,7,11,15-TetraMeC31 | 1.9 ± 0.5 | 0,82 | 2.03 ± 0.59 | 0,81 | 2.22 ± 0.46 | 0,95 |  |
| **3285** | **9-C33ene** | **0.31 ± 0.17** | **0,13** | **0.37 ± 0.14** | **0,15** | **1.59 ± 0.68** | **0,68** |  |
| **3293** | **7-C33ene** | **0.76 ± 0.32** | **0,33** | **0.74 ± 0.16** | **0,30** | **3.27 ± 1.2** | **1,41** |  |
| 300 | n-C33 | 1.02 ± 0.32 | 0,44 | 1.17 ± 0.35 | 0,47 | 1.32 ± 0.47 | 0,57 |  |
| 3334 | 9-; 11-; 13-; 15-MeC33 | 18.35 ± 5.89 | 7,96 | 22.55 ± 8.89 | 9,05 | 14.08 ± 4.07 | 6,06 |  |
| **3347** | **7-MeC33** | **1.52 ± 0.59** | **0,66** | **1.98 ± 0.72** | **0,79** | **0.1 ± 0.08** | **0,05** |  |
| **3353** | **5-MeC33** | **9.59 ± 3.36** | **4,16** | **9.78 ± 3.83** | **3,92** | **3.83 ± 0.75** | **1,65** |  |
| **3364** | **11,15-; 11,17-; 11,19-; 11,21-; 11,23-; 11,27-; 13,15-; 13,17-; 13,19-; 13,21-; 13,23-; 13,27-DiMeC33** | **10.25 ± 3.96** | **4,44** | **10.8 ± 4.1** | **4,34** | **2.95 ± 0.73** | **1,27** |  |
| **3371** | **7,15-; 7,17-; 7,19-; 7,21-; 23-DiMeC33** | **4.38 ± 1.54** | **1,90** | **4.63 ± 1.9** | **1,86** | **1.09 ± 0.38** | **0,47** |  |
| **3382** | **5,9-; 5,11-; 5,15-; 5,17-; 5,21-; 5,23-DiMeC33** | **7.36 ± 2.85** | **3,19** | **7.64 ± 2.24** | **3,06** | **32.68 ± 11.77** | **14,08** |  |
| **3384** | **11,13,21-; 11,15,21-; 11,17,21-; 11,13,23-; 11,15,23-; 11,17,23-TriMeC33** | **3.55 ± 1.48** | **1,54** | **3 ± 1.07** | **1,20** | **1.03 ± 0.2** | **0,44** |  |
| **3394** | **3,9-; 3,11-; 3,13-; 3,15-; 3,17-DiMeC33** | **3.3 ± 0.84** | **1,43** | **3.2 ± 0.7** | **1,28** | **6.89 ± 2.12** | **2,97** |  |
| 3427 | 5,9,13-; 5,11,13-TriMeC33 | 1.99 ± 0.54 | 0,86 | 2.18 ± 0.74 | 0,87 | 2.31 ± 0.58 | 0,99 |  |
| **3431** | **3,7,11-; 3,7,15-; 3,7,17-; 3,7,19-; 3,7,21-TriMeC33** | 0.89 ± 0.46 | 0,39 | **1.24 ± 0.73** | **0,50** | **0.44 ± 0.38** | **0,19** |  |
| 3458 | 3,7,11,15-TetraMeC33 | 3.94 ± 0.96 | 1,71 | 4.03 ± 1.16 | 1,62 | 3.3 ± 0.6 | 1,42 |  |
| **3477** | **5,13-; 5,15-; 5,17-; 5,19-DiMeC34** | **0.74 ± 0.23** | **0,32** | **0.71 ± 0.16** | **0,28** | **1.44 ± 0.39** | **0,62** |  |
| 3486 | 4,10-; 4,16-; 4,18-DiMeC34 | 0.18 ± 0.15 | 0,08 | 0.11 ± 0.12 | 0,04 | 0.2 ± 0.19 | 0,09 |  |
| 3531 | 11-; 13-; 15-; 17-MeC35 | 9.74 ± 2.84 | 4,22 | 12.15 ± 4.63 | 4,88 | 7 ± 1.66 | 3,01 |  |
| **3557** | **11,17-; 13,17-; 11,19-; 13,19-; 11,21-; 13,21-; 11,23-; 13,23-DiMeC35** | **26.66 ± 8.46** | **11,56** | **27.57 ± 10.24** | **11,06** | **8.36 ± 1.98** | **3,60** |  |
| **3568** | **7,19-; 7,21-; 7,23-DiMeC35** | **4.88 ± 1.4** | **2,12** | **4.48 ± 1.91** | **1,80** | **0.56 ± 0.33** | **0,24** |  |
| **3579** | **5,15-; 5,17-; 5,21-; 5,23-; 5,25-DiMeC35** | **9.25 ± 3.04** | **4,01** | **8.9 ± 3.11** | **3,57** | **29.75 ± 10.82** | **12,81** |  |
| **3591** | **3,17-; 3,19-; 3,21-; 3,23-; 3,25-DiMeC35** | **1.13 ± 0.48** | **0,49** | 1.02 ± 0.49 | 0,41 | **0.56 ± 0.24** | **0,24** |  |
| **3599** | **5,9,13-; 5,9,15-; 5,9,17-; 5,9,19-; 5,9,21-TriMeC35** | **1.66 ± 0.56** | **0,72** | **1.46 ± 0.42** | **0,58** | **3.84 ± 1.37** | **1,65** |  |
| **3648** | **3,13,19-; 3,13,23-; 3,13,25-; 3,17,19-; 3,17,23-; 3,17,25-; 3,19,23-; 3,19,25-; 3,23,25-TriMeC35** | **2 ± 0.62** | **0,87** | **1.98 ± 0.71** | **0,79** | **1 ± 0.23** | **0,43** |  |
| 3722 | 13-; 15-; 17-; 19-MeC37 | 1.81 ± 0.59 | 0,79 | 2.31 ± 0.94 | 0,93 | 1.52 ± 0.44 | 0,65 |  |
| **3744** | **11,17-; 11,19-; 11,21-; 11,23-; 13,17-; 13,19-; 13,21-; 13,23-DiMeC37** | **11.62 ± 3.61** | **5,04** | **11.79 ± 4.74** | **4,73** | **3.03 ± 0.88** | **1,30** |  |
| **3757** | **7,21-; 7,23-; 7,25-DiMeC37** | **1.55 ± 0.54** | **0,67** | **1.38 ± 0.63** | **0,56** | **0.16 ± 0.09** | **0,07** |  |
| **3767** | **5,15-; 5,17-; 5,23-; 5,25-; 5,27-DiMeC37** | **3.11 ± 1.01** | **1,35** | **2.96 ± 0.92** | **1,19** | **8.77 ± 3.22** | **3,78** |  |
